# Measuring gas vesicle dimensions by electron microscopy

**DOI:** 10.1101/2021.01.22.427725

**Authors:** Przemysław Dutka, Dina Malounda, Lauren Ann Metskas, Songye Chen, Robert C. Hurt, George J. Lu, Grant J. Jensen, Mikhail G. Shapiro

**Author notes:** Correspondence should be addressed to G.J.J. or M.G.S.

## Abstract

Gas vesicles (GVs) are cylindrical or spindle-shaped protein nanostructures filled with air and used for flotation by various cyanobacteria, heterotrophic bacteria, and Archaea. Recently, GVs have gained interest in biotechnology applications due to their ability to serve as imaging agents and actuators for ultrasound, magnetic resonance and several optical techniques. The diameter of GVs is a crucial parameter contributing to their mechanical stability, buoyancy function and evolution in host cells, as well as their properties in imaging applications. Despite its importance, reported diameters for the same types of GV differ depending on the method used for its assessment. Here, we provide an explanation for these discrepancies and utilize electron microscopy (EM) techniques to accurately estimate the diameter of the most commonly studied types of GVs. We show that during air drying on the EM grid, GVs flatten, leading to a ~1.5-fold increase in their apparent diameter. We demonstrate that GVs’ diameter can be accurately determined by direct measurements from cryo-EM samples or alternatively indirectly derived from widths of flat collapsed and negatively stained GVs. Our findings help explain the inconsistency in previously reported data and provide accurate methods to measure GV dimensions.

## INTRODUCTION

Gas Vesicles (GVs) are hollow, gas-filled protein nanostructures natively expressed in certain types of cyanobacteria, heterotrophic bacteria, and Archaea as a buoyancy aid (Walsby 1994). Recently, it was discovered that the unique physical properties of GVs enable them to serve as genetically encodable contrast agents for ultrasound and other imaging methods, allowing deep tissue imaging of cellular function (Shapiro, Goodwill, et al. 2014; Shapiro, Ramirez, et al. 2014; Bourdeau et al. 2018; Lu et al. 2018; Farhadi et al. 2019, 2020; Lakshmanan et al. 2020). In addition, GVs are being applied to acoustic manipulation and therapeutic uses of engineered cells (Bar-Zion et al. 2019; Wu et al. 2019).

Fully formed GVs adopt two predominant shapes - cylinders with conical ends or spindle-like. The GVs may be 0.1-2 μm in length, or even longer when heterologously expressed in more spacious mammalian cells (Farhadi et al. 2019). The mean diameter of GVs isolated from different species widely varies, but is relatively constant for the same type of GV. There is an inverse correlation between diameter and critical collapse pressure (Hayes and Walsby 1986). This correlation has important evolutionary consequences. While wider GVs can provide buoyancy at a lower energetic cost, they collapse at lower pressure. This is perhaps best reflected by analyzing the widths and collapse pressure of GVs isolated from *Planctronix spp.* from nordic lakes of different depths (Beard et al. 1999, 2000). Three types of GVs isolated from *Planctronix spp.* had widths of approximately 51, 58, and 67 nm with respective collapse pressures of 1.1, 0.9, and 0.7 MPa, allowing them to adapt to the hydrostatic pressure in different lakes (Beard et al. 2000; Dunton and Walsby 2005).

Despite the importance of GVs’ diameter for their biophysical properties, there are significant discrepancies in values reported in the literature. For example, the width of GVs from *Anabaena flos-aquae* (Ana) measured inside cells by thin-section electron microscopy (EM) was approximately 70 nm (Walsby 1971), which is considerably smaller than the value obtained by negative stain EM (ns-EM) for isolated GVs - 136 nm (Lakshmanan et al. 2017). Similar discrepancies can be observed for GVs from *Halobacterium salinarum* (Halo), whose reported values range from 45 nm to 250 nm (Simon 1981; Offner et al. 1998; Pfeifer 2012; Lakshmanan et al. 2017). To some extent, these discrepancies could be explained by natural variability in diameter. However, analysis of width distributions for GVs from several species of Cyanobacteria (Hayes and Walsby 1986) or *Bacillus megaterium* (Mega) (Farhadi et al. 2018) shows a narrow range. This inconsistency in diameter measurement was investigated almost 50 years ago by Walsby (Walsby 1971). He observed that Ana GVs have a constant width of 70 nm when measured inside cells by thin-section EM, which was close to the value measured for the purified sample imaged using a freeze-etching technique (75 nm). In contrast, estimations by ns-EM ranged from 70 to 114 nm (Walsby 1971). He suggested that the stain used in EM leads to swelling of GVs, which increases their diameter but has little effect on the length. As an alternative approach for assessing GV diameter, Walsby proposed indirect measurement based on the widths of flat collapsed GVs. The diameter of Ana GVs measured using this strategy was approximately 85 nm (Walsby and Bleything 1988). Archer and King gave another potential explanation for discrepancies in GV measurements. They proposed that the isolation process leads to deformations, increasing the width of GVs (Archer and King 1984). Regardless of these concerns, the diameter of GVs has been routinely assessed for isolated specimens by ns-EM.

As GVs have attracted more attention in biotechnology applications, accurate estimates of their diameter have become a critical input into GV engineering. For that reason, we investigated the discrepancies in reported GV diameters using modern microscopy tools. Using these updated techniques, we provide measurements for the most commonly studied GVs: Ana, Mega, and Halo. For Halo, we analyzed two different GV types, which are products of the independent gene clusters p-vac and c-vac.

## RESULTS AND DISCUSSION

To more closely evaluate the behavior of stained and air-dried GVs on the EM grid, we collected projection images for different types of GVs at 0° and 50° tilt and analyzed their morphology **(Figure 1a and 1b)**. Although we predicted some degree of distortions to the cylindrical shape of GVs, the observed differences were unexpectedly large. For Ana GVs, there was an average of 55 nm width difference between measurements at these two angles. The pattern was similar for both Mega and Halo GVs, although to a different degree. This data indicates that all types of GVs flatten during the staining procedure, adopting an elliptic cylinder shape.

**Figure 1.**
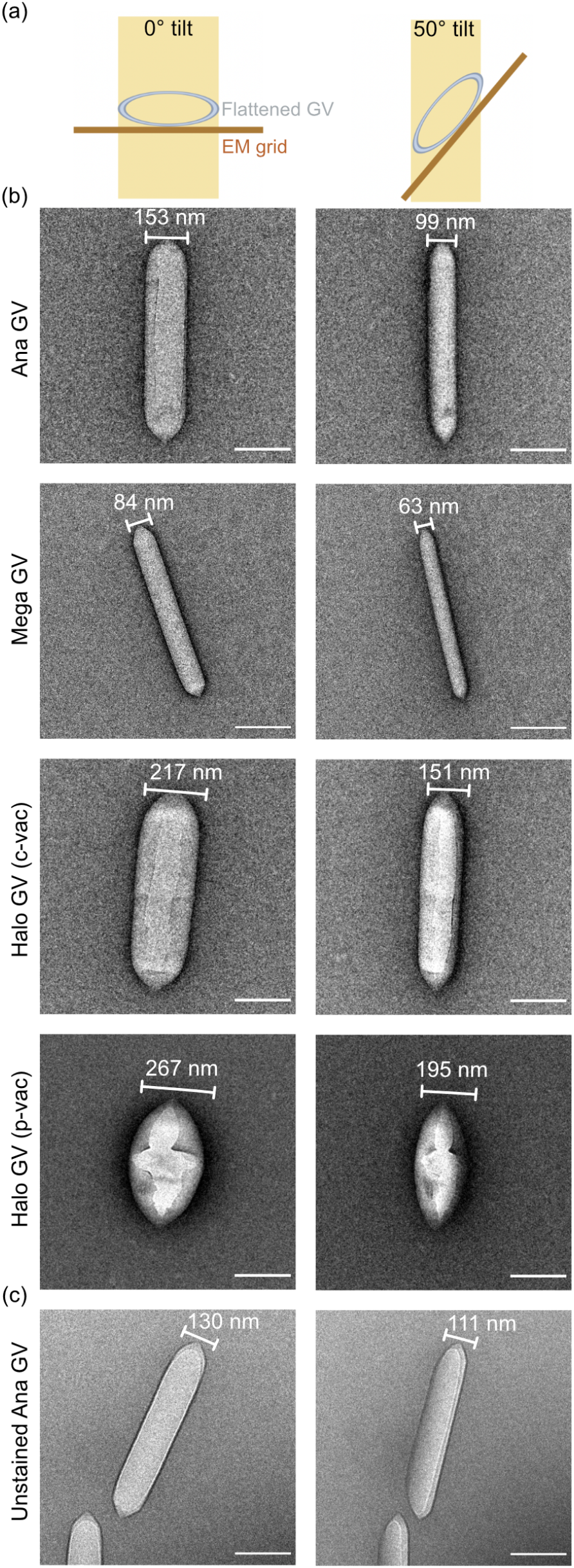
GVs flattening on the EM grid. (a) Schematic showing cross-section of the flattened GV at 0° and 50° tilt. (a,b) Representative projection images at 0° and 50° tilt for (b) negatively stained and air-dried Ana, Mega, and Halo GV; and (c) unstained, air-dried Ana GV. Scale bar, 200 nm.

Certain limitations of the ns-EM technology, such as specimen flattening or stain thickness irreproducibility, were previously described (Frank 2006). However, the observed deformation of the GV protein shell is not like the typical flattening reported before, where sample was mainly affected in z-direction with little to no effect on the x.y-dimensions (Frank 2006). Since GVs produce strong contrast on EM even without staining, we decided to take advantage of this unique property and evaluate the effect of the stain. Analysis of unstained, air-dried Ana GVs samples at 0° and 50° tilts show on average 20 nm difference in diameter **(Figure 1c)**, which is significantly less than the stained sample, but not negligible.

Distortions to the GV shape are the effect of the unique mechanical properties of GVs’ protein shell. In ns-EM, the sample lies on a carbon support; thus, we suspect that GVs are compressed by the surface tension of evaporating water. Notably, the degree of deformation appears to be correlated with critical collapse pressure. Halo GVs, which experience the most flattening, are also the least robust among investigated GVs, with collapse pressure of 0.1 MPa (Lakshmanan et al. 2017). In contrast, Mega GVs, which have a much higher collapse pressure of 0.7 MPa (Lakshmanan et al. 2017), flatten the least.

To obtain more accurate measurements of GV diameter, we used two complementary methods. First, we imaged the GVs with cryo-EM, which preserves GVs’ cylindrical shape. Unfortunately, cryo-EM is a more demanding technique, requiring time-consuming sample optimization, larger sample quantities, and access to a more sophisticated instrument. Alternatively, we inferred GV diameter from the widths of flat collapsed GVs with negative staining, as measured by Walsby and Bleything (Walsby and Bleything 1988). This method, which equates the collapsed GV width with half of the intact cylindrical circumference, should allow for a faster and more accessible estimation of GV dimensions. We decided to analyze diameter distribution for Mega, Ana, and Halo GVs using both strategies. Cryo-EM of intact GVs and collapsed ns-GV imaging resulted in similar values for each analyzed GV type **(Figure 2, Table 1)**, with differences within statistical error. Mega and Ana GVs appear to have a uniform diameter, varying within a narrow range **(Figure 2c, Table 1)**. In contrast, Halo GV diameters varied. Halo is capable of producing two types of GVs. Spindle-shaped GVs are encoded by the p-vac gene cluster located on an endogenous plasmid, while the c-vac cluster located on a mini-chromosome generates cylindrical GVs (Pfeifer 2012). According to our measurements, the diameter of both types of Halo GVs varies **(Figure 2c)**. However, some of this variability may be due to imperfect classification. All GV types begin their assembly as bicones, which look like smaller spindle-shape p-vac Halo GVs (Pfeifer 2012). Thus, some c-vac GVs, in their bicone phase, could have been classified as p-vac GVs. This misclassification could have made a minor contribution to the overall diameter distribution. Overall, the range of diameter values for different GV types suggest that Ana and Mega GVs have tighter regulation over diameter compared to Halo GVs. However, it is not yet known what the physiological consequences of this regulation are or how exactly the diameter is adjusted in growing GVs.

**Table 1.** Measured diameters (mean ± s.d.) for Mega, Ana, and Halo GVs by EM

| GV type | Intact GVs<br>(ns-EM)* (nm) | Intact GVs<br>(cryo-EM)<br>(nm) | Collapsed GVs<br>(ns-EM) (nm) |
| --- | --- | --- | --- |
| Mega | 73 ± 14 | 52 ± 6 | 54 ± 5 |
| Ana | 136 ± 21 | 85 ± 4 | 89 ± 6 |
| Halo (c-vac) | 251 ± 51 | 111 ± 32 | 116 ± 21 |
| Halo (p-vac) | | 182 ± 22 | 171 ± 19 |

\*Previously reported by Lakshmanan et al. (2017).

**Figure 2.**
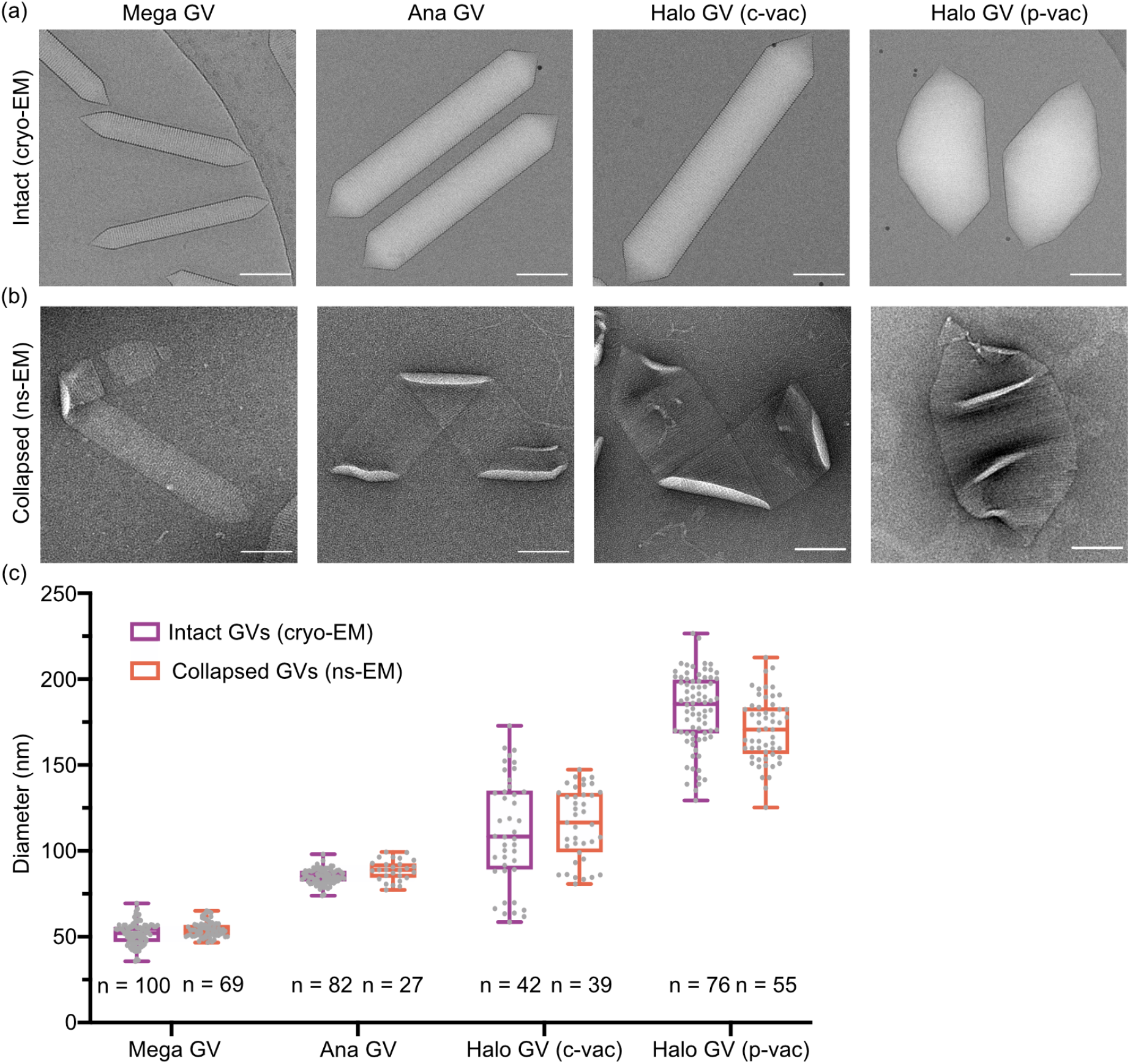
Diameter measurement for Mega, Ana, and Halo GVs. (a) Representative cryo-EM of intact GVs used for direct diameter measurement. (b) Representative ns-EM images of collapsed GVs used for indirect diameter assessment based on widths of flat collapsed regions. Scale bar, 100 nm. (c) Diameter distribution for Mega, Ana, and Halo GVs measured by cryo-EM and collapsed ns-EM. Center line indicates median, the box limits denote the interquartile range and the whiskers absolute range. Each dot represents an individual measurement.

Taken together, our findings provide an explanation for discrepancies in previous GV diameter measurements reported in the literature. Although ns-EM is routinely used to evaluate the morphology and dimensions of intact GVs (Li and Cannon 1998; Ramsay et al. 2011; Xu et al. 2014; Lakshmanan et al. 2017; Farhadi et al. 2018), our data show that this method causes GV flattening and inaccurate apparent diameter. Instead, cryo-EM of intact GVs and ns-EM of flat collapsed GVs provide correct dimensions that are mutually consistent between the two methods, as shown here for three commonly studied GVs variants.

## MATERIALS AND METHODS

### GV expression and purification

GVs were either isolated from native sources (Ana and Halo) or expressed heterologously in Rosetta 2(DE3) pLysS *E.coli* cells (Mega) as previously described (Lakshmanan et al. 2017). In the final two or three rounds of buoyancy purification, sample buffer was exchanged to 10 mM HEPES, pH 7.5. Concentrations were measured by optical density (OD) at 500 nm using a spectrophotometer (NanoDrop ND-1000, Thermo Scientific).

### Negative stain electron microscopy

For imaging of intact GVs, the purified sample was diluted to OD_500_ ~0.5 for Ana and Halo, and OD_500_ ~0.2 for Mega. When data for collapsed GVs were collected, diluted samples were squeezed in a sealed syringe until turned transparent. 3 μL of the target sample was applied to a freshly glow-discharged (Pelco EasiGlow, 15mA, 1 min) Formvar/carboncoated, 200 mesh copper grid (Ted Pella) for 1 min before blotting. Afterward, the sample was incubated for 1 min with a 0.75% uranyl formate solution before blotting and air-dried. Image acquisition was performed using a Tecnai T12 (FEI, now Thermo Fisher Scientific) electron microscope at 120 kV, equipped with a Gatan Ultrascan 2k X 2k CCD.

### Cryo-electron microscopy

For cryo-EM, Quantifoil R2/2 200 Mesh, extra thick carbon, copper grids (EMS) were glow discharged (Pelco EasiGlow, 10mA, 1 min). Freshly purified Mega (OD_500_ ~1), Ana (OD_500_ ~15), and Halo (OD_500_ ~8) GVs sample was frozen using a Mark IV Vitrobot (FEI, now Thermo Fisher Scientific) (4°C, 100% humidity, blot force 3, blot time 4s). Micrographs were collected on a 300kV Titan Krios microscope (FEI, now Thermo Fisher Scientific) with an energy filter (Gatan) and equipped with a K3 6k x 4k direct electron detector (Gatan). Data were collected using SerialEM software with a pixel size of either 1.4 Å (x64,000 magnification) or 2.15Å (x42,000 magnification) and −2.5 μm defocus (Mastronarde 2005).

### Diameter measurement

All measurements were made using IMOD software (Kremer, Mastronarde, and McIntosh 1996). The cylinder/spindle diameter direct measurements from cryo-EM micrographs were performed only once for each GVs at its widest region. Indirectly diameter was calculated as 2*w/π*, where *w* is the width of the flat collapsed gas vesicle measured from the ns-EM micrograph. Sample from at least two independent preparations were used for each measurement.

## ACKNOWLEDGEMENTS

This work was supported by the National Institutes of Health (grant R35-GM122588 to G.J.J. and R01-EB018975 to M.G.S.) and the Caltech Center for Environmental Microbial Interactions (CEMI). Electron microscopy was performed in the Beckman Institute Resource Center for Transmission Electron Microscopy at Caltech. Related research in the Shapiro Laboratory is also supported by the Heritage Medical Research Institute, the Pew Scholarship in the Biomedical Sciences, and the Packard Fellowship for Science and Engineering.

## AUTHOR CONTRIBUTIONS

**Przemysław Dutka:** Conceptualization; methodology; investigation; formal analysis; visualization; writing – original draft preparation; writing – review editing. **Dina Malounda:** Investigation. **Lauren Ann Metskas:** Investigation. **Songye Chen:** Investigation. **Robert C. Hurt:** Investigation. **George J. Lu:** Investigation. **Grant J. Jensen:** Conceptualization; writing – review editing; supervision; funding acquisition. **Mikhail G. Shapiro:** Conceptualization; writing – review editing; supervision; funding acquisition.

